# Motivation upregulates the adaptive response in sensorimotor learning

**DOI:** 10.1101/2023.09.17.558102

**Authors:** Salma Khatib, Vikram S. Chib, Firas Mawase

**Author notes:** **Correspondence to:** Firas Mawase. These authors contributed equally (Vikram S. Chib, Firas Mawase).

## Abstract

Motivational state plays a critical role in our ability to learn new motor skills; however, the mechanisms by which motivation influences motor learning are poorly understood. Using a motor learning paradigm in which motivation was varied in a trial-by-trial manner, we found that motivation affects learning by upregulating the rate of the adaptive response, increasing individuals’ speed of learning. This unveils previously unidentified evidence for a mechanism through which motivation shapes error-based motor learning.

## Main

Being in a heightened motivational state can drive more vigorous motor responses^1–5^, increase effortful exertion^6–8^ and the velocity of movement^9–11^, and reduce reaction times^12,13^ . These effects are ubiquitous, having been shown across various tasks, in rodents, nonhuman primates, and humans. Recent studies of motor learning have begun to examine how rewards, contingent on motor performance, influence adaptation^14,15^ and skill learning^12,16,17^. When successful motor performance was rewarded, motor learning was accelerated, supporting the idea that performance-contingent rewards motivate motor learning. However, these studies did not dissociate the influences of reward and motivation on learning, making it difficult to understand the distinct influence of motivational state on adaptation.

Previous experimental approaches motivated participants by rewarding the successful completion of a motor task during adaptation^14,15^. However, motivational state in these experiments was confounded by the performance-contingent outcomes -during the trial-by-trial motor adaptation process, participants not only learned from previous motor errors but also reward delivery, which was contingent on performance. These experimental designs made it difficult to dissociate the influence of motivation and reward learning on motor adaption.

To study how changes in motivational state influence adaptive responses during motor learning, we developed a behavioral paradigm in which participants were exposed to cues associated with different motivational states while interacting with environments that imparted sensorimotor perturbations during arm-reaching movements. Such sensorimotor perturbations have been used extensively to study motor learning, where a sensory prediction error (e.g., the difference between the predicted and perceived location of the hand) leads to updates of an internal model. To decouple the effect of motivation from performance-contingent outcomes during sensorimotor adaptations, and make these two components statistically independent, we stochastically presented participants with different motivational cues and sensorimotor perturbations. This made the motivational cues orthogonal to motor errors and learning performance, allowing us to isolate their influence on motor adaptation.

During experiments, participants were instructed to grasp the hand of a two-joint planar robotic manipulandum, and make rapid reaching movements with the dominant hand toward a single target (Fig. 1A). Participants first performed a baseline phase, in which no external force-field perturbations were applied (i.e., null-field trials), to familiarize them with the motor task. They were then exposed to pseudo-random velocity-dependent force-field perturbations *f*. The force that perturbed participants’ hands was orthogonal to their direction of hand movement and proportional to their movement speed. Across trials, the perturbation magnitude was uniformly distributed and sampled from the range [±30%, ±60%, ±90%] × *f*, in both the clockwise (CW, +) and counterclockwise (CCW, -) directions. Critically, some perturbation trials were associated with monetary cues that were uniformly sampled, but statistically independent from the perturbation schedule in the range [₪10,₪ 100]. These cues were used to set the motivational state to be low (₪10) or high (₪100) on each trial (Fig. 1B&C). Participants did not see the reward outcomes at the end of trials, to minimize the influence of reward delivery on motivation, and isolate the motivational aspects of reward.

**Figure 1.**
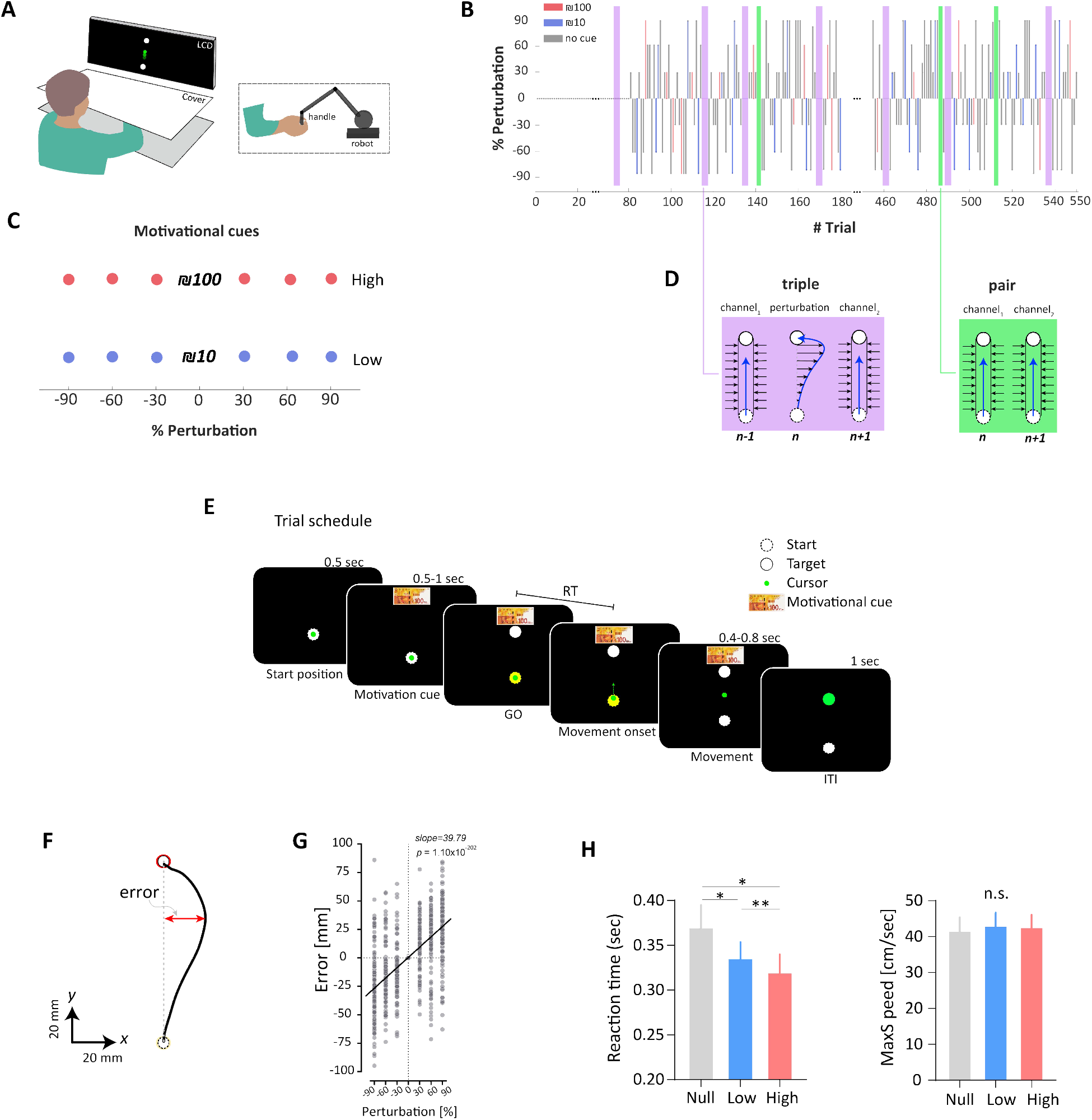
Experiment design, error quantification and the effect of motivation on performance. **A** Experimental setup. Participants grasped the handle of a robotic manipulandum and made planar reaching movement toward a single target located 90° from the start point. **B**. Participants performed a multi-phase experiment with a baseline block and adaptation block. The size of perturbation and motivational cue vary randomly across trials. Participants exposed to a random perturbation from the range of [±30%, ±60%, ±90%] of a predefined value velocity-dependent force-field perturbations ***f***. **C**. No statistical dependency between perturbation size and motivation cue during the adaptation phase (correlation ∼ 0). Three types of cues were presented: null (no motivation cue), low (₪10) and high motivation (₪100). **D**. To quantify adaptative response, we introduced triples (magenta) and pair (green) trials including force channels (error-clamped) to measure the force motor output. **E**. Time schedule of a trial. **F**. Movement trajectory in the perturbation trial. Movement error was calculated as the deviation of the actual trajectory at max speed from the straight line connecting the start point and the center of the target. **G**. Errors as a function of perturbation sizes pooled across all participants. **H**. Motivation influences the reaction time (left) but not the speed of the movement (right). Bars represent mean and error-bars represent s.e. of the mean. * for *p* < 0.05, ** for *p* < 0.01 and ***n.s***. for not-significant.

To quantify adaptation, we measured the motor output before and after each force-field perturbation trial. We pseudorandomly presented groups of three trials (i.e., triples) in which a force-field perturbation trial (P) was presented between force channel trials (C) (C_1_ → P → C_2_). During channel trials, participants made reaching movements through a virtual environment where stiff virtual walls were rendered that constrained the hand path to a straight line with minimal lateral deviation and no velocity-dependent force. We contrasted the forces participants exerted against the channel in *C*_1_ and *C*_2_ to generate a measure of adaptive response (Fig. 1D and Fig. S1). By doing so, we could measure the motor output in a specific trial in response to error in the previous trial. These triples of channel-perturbation-channel (*C*_1_*PC*_2_) trails were randomly separated by 0, 1, or 2 regular trials, preventing participants from predicting their timing^18,19^.

To evaluate how motivational state influences adaptation, participants were presented with three different types of triples *C*_1_*PC*_2_, 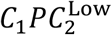, and 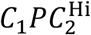. In *C*_1_*PC*_2_, no motivational cue was presented and, therefore, the change in force from *C*_1_ to *C*_2_ was due solely to the error experienced in trial *P*. In 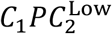, and 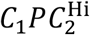 low and high motivational cues were presented in the second force channel trial, so the change in force from *C*_1_ to *C*_2_ was not only due to the error experienced in trial *P*, but also the motivational cue presented in *C*_2_.

To quantitatively characterize the adaptive response from force field perturbations and changes in motivational state, we considered a model that accounted for motor output across triples. Considering that in trial *n* −1 participants experience a channel trial *C*_1_, associated with a measured motor output *u*^*n*-1^, the motor output that the participant produces in the next two trials can be characterized by three factors: 1) a forgetting rate α, 2) the error that was experienced during the perturbation trial at time *n, e*^(*n*)^ (i.e., the difference between the predicted and observed feedback), and 3) the adaptive response from error, 𝜆(*e*^(*n*)^), as follows:

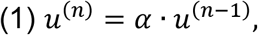

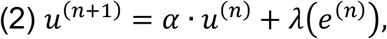

From Eq. (2), the change in motor output due to the trial-by-trial adaptive response to error, is:

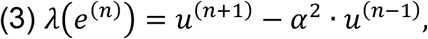

To estimate the motor output *u*^(*n*)^, we regressed the measured force *f*(*t*) that the participant had produced against the force channel during perturbation trials onto the ideal force 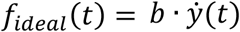 and found the parameters that minimized the quantity *f*(*t*) = *k*_1_ · *f*_*ideal*_(*t*) + *k*_0_, and then set *u*^(*n*)^ = *k*_1_.

To estimate the forgetting rate α, we introduced additional pairs of channel trials *C*_1_*C*_2_ (Fig. 1D) with no perturbation, and calculated the ratio of the forces in these two trials (averaged across all pairs):

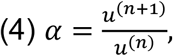

Using the computational framework above, we were able to estimate the adaptive response to null (*C*_1_*PC*_2_), low 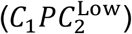 and high 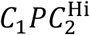 motivation triples separately.

Our data revealed a significant relationship between force field perturbation magnitude and direction, and movement error during perturbation trials, irrespective of the motivational cue, (regression slope of +39.79, *p* < 0.0001, Fig. 1F&G). In concert with previous motor adaptation studies, larger force field perturbations resulted in increased movement errors that were proportional in magnitude and direction to the perturbation^20^. Next, we examined the influence of motivational state on motor performance by examining reaction time and peak movement velocity (i.e., vigor). We found that reaction time decreased as the motivational salience of cues increased (*F*_1,13_ = 8.81, *p* < 0.0095), while peak velocity did not significantly differ across motivational conditions (*F*1,13 = 1.73, *p* = 0.20). These results show that while motivation influenced the speed of reaction time, as previously shown^13^, it did not influence motor execution speed (Fig. 1H).

To assess how motivational state modulates motor adaptation, we computed the amount learned from error (using triples) in a trial-by-trial fashion. We measured the change in motor output from the trial before and after the perturbation (i.e., the adaptive response). We found that adaptive response was linearly related to the amount of motor error experienced during perturbation trials, and the level of the motivational state modulated the extent of the adaptive response (Fig. 2). A higher motivational state was associated with increased adaptive responses, suggesting that motivation had a faciliatory effect on motor adaptation (*F*1,13 = 28.32, *p* < 0.0001). Essentially, a higher motivational state increased the rate at which changes in motor performance influenced learning from error (i.e., sensitivity, Fig. 2C). Post-hoc analysis revealed that sensitivity to errors in the High motivational state was significantly (*t*_13_ = 5.81, *p* = 0.0002, holm-šídák’s correction for multiple comparisons) larger than the sensitivity in Low motivation state, and the Null condition (*t*_13_ = 5.91, *p* < 0.0002).

**Figure 2.**
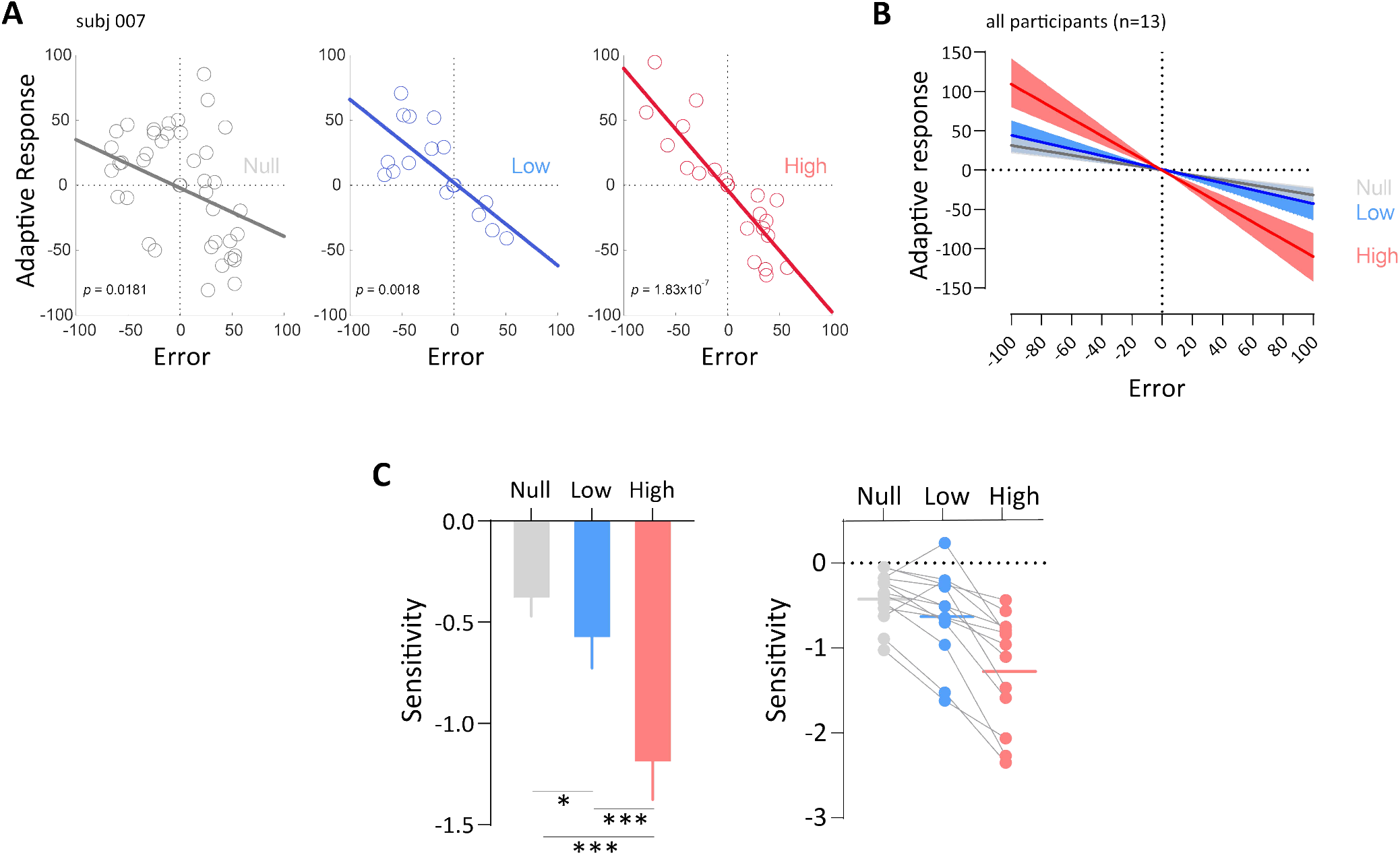
Motivation alters the rate of adaptive response. **A** Adaptive response as a function of the experienced error for one participant (#007). Left – trials with no motivation cues in the 2^nd^ force channel (α_2_), Middle – trials with low motivation cue in *C*_2_ and Right-trials with high motivation cue in *C*_2_. **B**. Adaptive response for all participants pooled together. Solid lines represent the regression line best fit the data and shaded area represent %95 confidence intervals (CI) of the fitted line. **C**. Motivation effected learning by increasing the rate (i.e., sensitivity) of the adaptive response. Left-group average data of sensitivity, in each of the motivation conditions. Right-individual data of the sensitivity. Bars represent mean and error-bars represent s.e. of the mean. * for *p* < 0.05, and *** for *p* < 0.001.

Our results show that a heightened motivational state upregulates the speed of motor learning by increasing the sensitivity to experienced error. Previous work has shown that positive reward-based feedback increased retention of motor memories, while negative punishment-based feedback, improved learning rate. However, these studies motivated participants by delivering rewards and punishments that were contingent on completion of the motor task^15,17,21^. These paradigms make it difficult to dissociate the influences of motivation from changes in expected value -at the beginning of adaptation, before learning, potential reward is near zero since participants have not yet learned the task; and the potential for reward is maximized and stable only when participants fully adapt to the task. Here we designed an adaptation paradigm to decouple the effect of motivation from the performance-contingent rewards by making these two components statistically independent. Specifically, we implemented a force-field perturbation paradigm that allowed for a dissociation of the adaptive response of each of the two components by fully randomizing external perturbations, and thus motor errors, and making the external motivational cues orthogonal to performance. Our paradigm allowed us to demonstrate a powerful, and hitherto undocumented, mechanistic understanding of the beneficial effect of motivation on sensorimotor learning, suggesting that motivation during learning is a crucial factor to consider when examining modulators of motor behavior acquisition.

One open question concerns how motivational state interacts with motor adaption at the neural level. While error-based motor learning highly depends on the cerebellum^22–26^, the basal ganglia is considered to be essential for motivation/reward-based (reinforcement) learning^27–29^. Recently the cerebellum has also been shown to be involved in motivation-based adjustment of motor output, a function that has more traditionally been associated with the basal ganglia. Anatomically, the cerebellum and the basal ganglia are interconnected at the subcortical level. The subthalamic nucleus in the basal ganglia sends dense disynaptic projections to the cerebellar cortex^30,31^. Similarly, the dentate nucleus in the cerebellum sends dense disynaptic projection to the striatum. These observations lead to a new functional perspective that the cerebellum and the basal ganglia form an integrated network that is essential for motivated motor performance and learning. In the future it will be important to explore the effect of motivation in sensorimotor learning as a byproduct of the complex interplay between the learning mechanisms supported by the basal ganglia and the cerebellum network.

## Methods

### Participants

13 right-handed participants (6 female), aged 25.8±3.7 years (mean±STD), were recruited for the current study. Participants provided written consent to participate in the study, which was approved by the Technion Institutional Review Board. All participants were deemed fully capable in terms of motor abilities, with no history of brain damage or motor impairments affecting hand/arm movements.

### Experimental Task

Participants were required to hold the handle of a robotic arm (1.5 Phantom) and make reaching movements (Fig. 1A) with the dominant hand to guide a cursor toward a signal target (15 mm diameter), which appeared 10 cm from a central start location (15 mm diameter). To ensure consistent movement speed and encourage participants to execute a single movement on each trial, participants were required to maintain their movement duration (the time to reach a displacement of ∼10 cm from the start location) between 400-800 msec. They were provided feedback in the form of a text appeared on the screen with “move faster!”, if they moved faster or “move slower!” if they moved slower than the criterion, on any given trial. A movement considered successful if the cursor ended within a predefined distance from the target and the cursor was brought into the target in time. The color of the target turned green when movement was successful. The predefined time window was selected to diminish opportunity for online corrections. The position and velocity of the robot’s handle were sampled at 200 Hz through a custom C++ (visual studio) script.

Participants performed two blocks of reaching movements. In the first block (i.e., baseline), participants performed 80 trials in which no external force-field perturbations were applied (i.e., null-field trials). In the second block (i.e., adaptation), participants performed 470 trials. In these trials, participants were exposed to pseudo-random velocity-dependent force-field perturbations (i.e., FF trials), in the form of 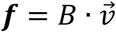, where 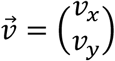 is the participant’s hand velocity and 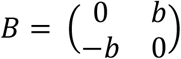 is the damping constant. We set *b* to be 13 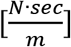, indicating 100% of the FF. Across trials, the perturbation magnitude was uniformly distributed and sampled from the range [±30%, ±60%, ±90%] × *f*. Positive sign of force implies clockwise (CW) perturbation whereas negative sign implies counterclockwise (CCW). Critically, within these perturbation trials, some trials were associated with monetary cues that are uniformly sampled, but statistically independent of the perturbation schedule, from the range [₪10,₪ 100]. These cues were used to set the motivational state to be high (₪100) or low (₪10) in each trial, and before the participants started the movement. To minimize the influence of reward delivery on motivation, participants did not see the reward outcomes at the end of trials. At the beginning of the experiment, participants were informed that they would receive a fixed show-up fee of (₪80=∼$22) and that this fee did not depend on performance or behavior during the experiment. They were also informed that a single motivational trial will be randomly selected by the end of the experiment and the monetary cue appeared in this selected trial will be given to them as an extra. That is, participants in our experiment can earn either 80+10=₪90 or 80+100=₪180.

### Time schedule of a trial

Each trial began after the cursor was held in the starting position for a 0.5 sec. After this and for each of the motivational trials, individuals were shown a motivational cue image for 0.5-1 sec (time changes randomly across trials within this range). On trials where no motivational cue is used, no cue images were presented, and the screen remained blank for the same amount of time as in the motivational trials. Then, the target appeared at a fixed location and the color of the start point turned yellow. The appearance of the target and the change of the color served as a Go cue to start the reaching movement. The motivational cue appeared during the movement and disappeared as soon as participants got close to the end of the movement. Feedback of cursor position was given continuously during the movement. By the end of the movement, the target turned green if participants reach the target in the allowed time (0.4-0.8 sec) and distance, and red otherwise. Inter-trial-interval (ITI) was set to be 1 sec and participants were asked to avoid any movements during this time (Fig. 1E).

## Data analysis

Kinematic data were collected from the robot at 200 Hz and stored on a computer for off-line analysis using MATLAB (The MathWorks, Natick, MA). Movement performance was quantified at each trial (null channel) by calculating the end point error (mm), defined as the distance between the movement completion location and the target location. Positive values indicated clockwise errors whereas negative values indicated counterclockwise errors.

### Measuring motor output (i.e., u^(n)^) in force channel trials

Throughout the experiment, we introduced force channel at different trials. Force channel trial (i.e., *C*) were similar to other trials in the sense that the participants did not receive different instructions; however, in these trials, the haptic device constrained participants’ movement by enclosing the straight path between the center of the cursor at trial initiation and the end location within high-stiffness virtual walls. The virtual walls were implemented by applying a one-dimensional spring force (spring coefficient k=2.5 k·N/m) and a damping force (damping coefficient b=25 N·s/m) around the channel. These force channels allowed us not only to minimize lateral deviation (i.e., minimizing sensory prediction error) but also to measure lateral forces that the participant applied during the reach. The rationale for this paradigm was that if participants have an internal representation of the perturbing forces, it should be reflected in the forces that they apply on the force channel as a mirrored profile of the representation of the perturbation^32–34^. Finally, to estimate the motor output *u*^(*n*)^, we first regressed the measured force *f*(*t*) that the participant had produced against the force channel onto the ideal force 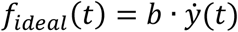 and then found the parameters that minimized the quantity *f*(*t*) = *k*_1_ · *f*_*ideal*_(*t*) + *k*_0_, and then set *u*^(*n*)^ = *k*_1_.

### Quantification of the adaptive response to errors (𝜆(e^(n)^)

We used a computational state-space model implemented in triples *C*_1_*PC*_2_ to estimate the adaptive response. Briefly, suppose that in trial *n* -1 we have a channel trial *C*_1_, where we measure motor output *u*^*n*-1^. The motor output that the participant produces in the next two trials can then be characterized by three factors: 1) a forgetting rate α, 2) error that was experienced in trial *n*, labeled as *e*^(*n*)^ (i.e., the difference between the predicted and observed feedback), and 3) adaptive response from error, as follows: 𝜆(*e*(^*n*)^) = *u*(^*n*+1)^ -α^2^ · *u*(^*n*-1)^. To estimate the forgetting rate α, we introduced additional pairs of channel trials *C*_1_*C*_2_ (Fig. 1B&D) with no perturbation and calculated the ratio of the forces in these two trials (averaged across all pairs) as follows α = α^(*n*+1)^α^(*n*)^.

We dissociated the adaptive response to errors from a response to motivational state, by introducing two types of triples,*C*_1_*PC* and 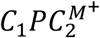 . The main difference between these triples is that while in *C*_1_*PC*_2_, no motivational cues was presented, in 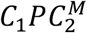, a motivational cue was presented in the second force channel trial so the change in force from *C*_1_ to *C*_2_ is not only due to the error experienced in trial *P* that occurred between *C*_1_ and *C*_2_, but also due to the motivational cue presented in *C*_2_.

### Sensitivity to errors

We measured the sensitivity to errors as the ratio between change in the adaptive response and the change in errors (i.e., slope of the adaptive response), as follows

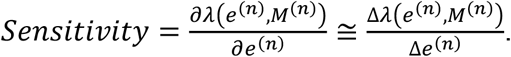

#### Reaction time (RT)

We calculated RT as the time interval between the time of stimulus presentation (i.e., target) and the time of movement onset (defined as time at 5% of max velocity).

## Acknowledgments

This was supported by the United States -Israel Binational Science Foundation (BSF) Grant 2021323 (FM and VSC).

## Supplementary Information

**Fig. S1.**
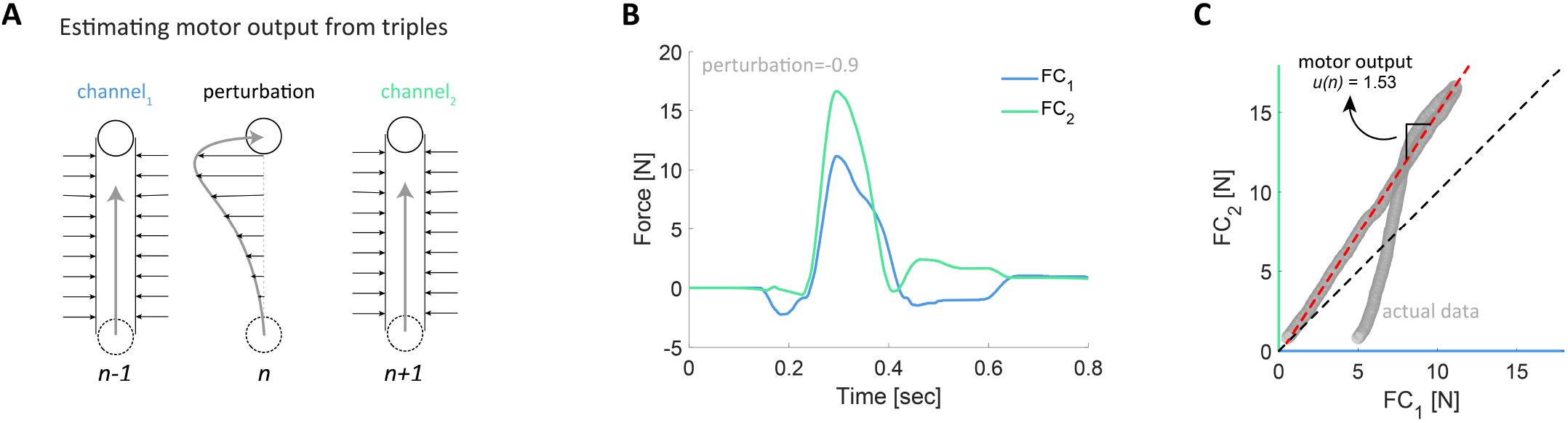
Estimating the motor output (α(^*n*)^) from *C*_1_*PC*_2_ triples. **A**. In order to estimate the adapted motor output in response to an experienced error, we have implemented triples, in different locations throughout the experiment, that consisted of a force channel trial, regular trial and then a second force channel. **B**. Force profile as a function of time in the first and second force channels following a perturbation of 90% counterclockwise force field. Data from an individual participant. **C**. A regression line was fitted to the force data in ***C***_2_ as a function of force data in ***C***_1_. Motor output (α(^*n*)^) was then calculated as the slope of the regression line best fitted the data.

**Fig. S2.**
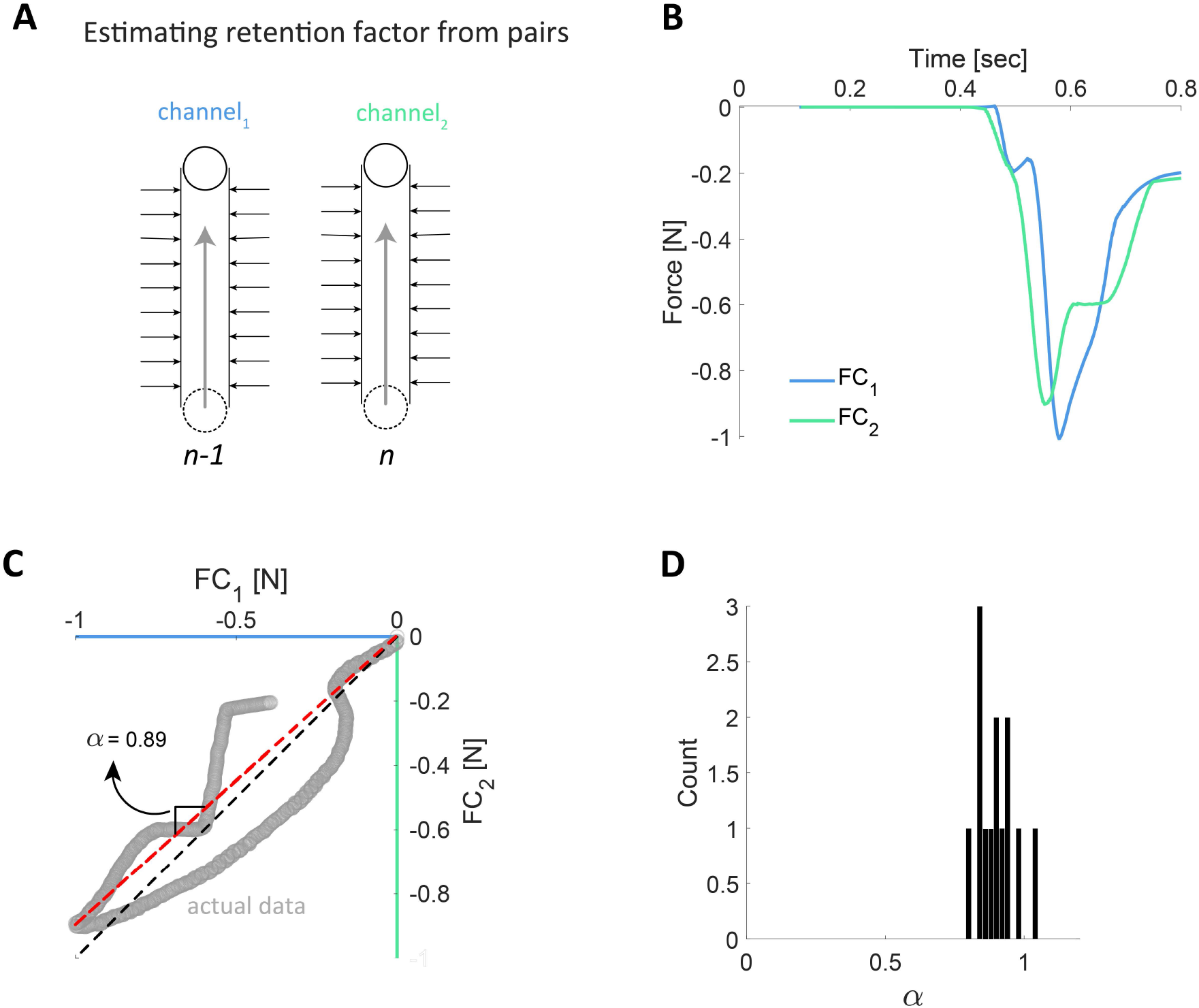
Estimating the retention factor (α) from *C*_1_*C*_2_ couples. **A**. We used consecutive force channel trials to estimate the retention (i.e., decay) of the adapted state from one trial to the next trial, in the absence of experienced error. **B**. Force profile as a function of time in the first and second force channels. Data from an individual participant. **C**. A regression line was fitted to the force data in ***C***_2_ as a function of force data in ***C***_1_. Retention factor (α) was then calculated as the slope of the regression line best fitted the data. **D**. Distribution of α across participants.

**Fig. S3.**
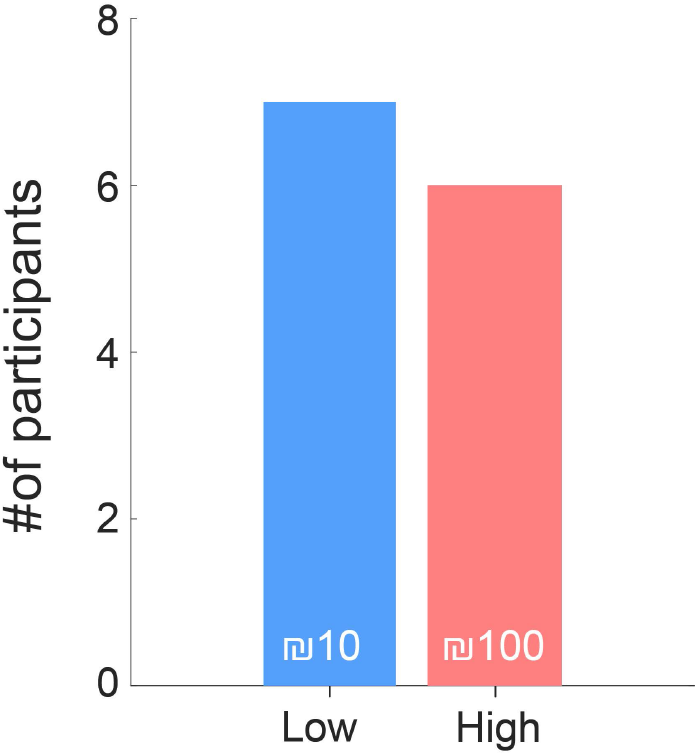
Distribution of participants by final compensation. Confirmatory analysis indicates no difference between the number of participants receiving high (n=6) and low (n=7) compensation at the conclusion of the experiment.

